# Effects of aging on semantic and episodic contributions to false memory

**DOI:** 10.64898/2026.08.21.746257

**Authors:** Isabelle L. Moore, Nicole M. Long

## Abstract

Healthy older adults are more susceptible to false memories than young adults. Traditional false memory paradigms leverage semantic overlap, shared meaning, to induce false memories, but experiences can also overlap temporally whereby they occur close together in time. Prior work shows that older adults have impaired episodic memory, memory for events within a spatiotemporal context, corresponding to an overall shift toward semantic memory and away from episodic memory across the lifespan. We hypothesize that compared to young adults, older adults rely more heavily on semantic versus episodic information, which promotes false memory. We collected behavioral data in young and older adults performing an old/new recognition memory task in which we manipulated the degree of semantic and temporal overlap between study words and included critical lures, unstudied words that semantically overlap with study words. We find that whereas young and older adults are similarly reliant on semantic relative to episodic information to support false memory, the two age groups differ in their reliance on semantic relative to episodic information to support true memory. These results suggest that differences in the orientation of attention – toward semantic vs. episodic information – may underlie age-related memory changes.

**Public Significance Statement:** The present findings demonstrate that young and older adults differ in their use of gist-based (semantic) information versus event-specific (episodic) information in true memory but not false memory. These results suggest that age-related memory changes may arise from differences in the orientation of attention toward the semantic or episodic dimensions of events.

## Introduction

Healthy aging is associated with increases in false memory (Koutstaal & Schacter, 1997), remembering events differently than how they happened (Roediger & McDermott, 1995). For example, having seen van Gogh’s *Sunflowers*, an older adult may be more likely than a young adult to falsely remember having seen van Gogh’s *Almond Blossoms*. Along with increased susceptibility to false memory, older adults have selective deficits in episodic memory, memory for events within a spatiotemporal context (Tulving, 1972). In contrast, semantic memory, memory for facts and concepts, is typically intact in healthy aging (Wingfield & Kahana, 2002). An older adult may therefore be able to remember general facts about the artist’s life (e.g. *van Gogh lived in France*), but have difficulty remembering the specific details of when and where they viewed one of his paintings (e.g. *last month at the Metropolitan Museum of Art*). Memory for episodic and semantic details are negatively correlated, an effect which increases with age (Devitt, Addis, & Schacter, 2017), and corresponds to an overall shift toward semantic memory over the lifespan (Ofen & Shing, 2013). Over-reliance on semantic memory may underlie older adults’ increased susceptibility to false memories (Koutstaal, Schacter, & Brenner, 2001), as older adults are more likely to experience false memories when those memories are consistent with prior knowledge (Norman & Schacter, 1997). The aim of the present study is to investigate the extent to which age-related increases in false memory arise from differential reliance on semantic versus episodic processing.

Older adults may rely on semantic memory to compensate for diminished episodic memory. Relative to young adults, older adults show memory impairments for episodic associations (Wilkniss, Jones, Korol, Gold, & Manning, 1997) – connections between events that occur close in time or space. Older adults also show decreased temporal organization during free recall tasks, suggesting a diminished ability to encode and/or retrieve temporal information (Wingfield, Lindfield, & Kahana, 1998). However, older adults do not differ from young adults in memory for items (Wilkniss et al., 1997; De Brigard, Langella, Stanley, Castel, & Giovanello, 2019) or semantic associations (Amer, Giovanello, Nichol, Hasher, & Grady, 2019) – connections between events that share meaning. Memory for episodic and semantic details are negatively correlated such that the more information individuals generate about generalized event knowledge and personal facts devoid of context (semantic information), the less information they generate about time and place (episodic information), an effect which increases with age (Devitt et al., 2017). This corresponds to an overall shift toward semantic memory that occurs across the lifespan (Ofen & Shing, 2013). Furthermore, prior knowledge may be deployed to counteract insufficient episodic representations in memory. When studying realistic and unrealistic prices for grocery store items, older adults’ memory performance is comparable to young adults only for those items that are priced realistically, suggesting that older adults utilize prior knowledge to ameliorate performance (Castel, 2005; Amer, Giovanello, Grady, & Hasher, 2018). Together, these findings suggest that older adults may have a semantic bias – a shift toward attending to semantic information when faced with a paucity of episodic information.

Though potentially compensatory, the shift in attention to semantic information may increase susceptibility to false memories in older adults. Young adults are more likely to experience false memories when study items semantically overlap (Gutchess & Schacter, 2012). False memories are often studied using the Deese-Roediger-McDermott (DRM) paradigm in which participants study multiple words that overlap semantically and are tested on both those items as well as critical lures, unstudied items that semantically overlap with study items (Roediger & McDermott, 1995). Older adults are more likely to falsely remember critical lures than young adults in the DRM paradigm (Norman & Schacter, 1997), and both young and older adults are more likely to falsely remember critical lures as more semantically overlapping items are presented during study (Koutstaal & Schacter, 1997). Thus, semantic processing can negatively impact memory by increasing the tendency to false alarm. An age-related bias toward semantic information may therefore account for older adults’ increased false alarm rates.

Episodic information, such as the temporal features of events, additionally has the potential to impact false memory susceptibility. Temporal features are thought to be contained in or a by-product of a slowly drifting context representation, an amalgamation of both external stimuli and internal states (Estes, 1955; Bower, 1972; Howard & Kahana, 2002). Context is bound to the present experience (Wang & Diana, 2017; Long & Kahana, 2019; Yonelinas, Ranganath, Ekstrom, & Wiltgen, 2019) and cues retrieval (Long et al., 2017). Context includes a mix of semantic and temporal features and drifts over time, such that events that share meaning and/or occur close together in time are bound to similar context representations (Polyn, Norman, & Kahana, 2009; Long & Kahana, 2017). Young adults can leverage temporal features to support memory, as evidenced by young adults’ tendency to consecutively recall items presented in neighboring serial positions on study lists in free recall (Kahana, 1996; Sederberg, Miller, Howard, & Kahana, 2010). In addition to facilitating true memory, access to temporal features may also enable resistance to false memory in young adults. Given that study items in the DRM paradigm share a temporal context – the study list – whereas critical lures do not, temporal features could potentially be used to enable rejection of critical lures. According to this account, temporal overlap between study items would *decrease* false memory susceptibility. Alternatively, temporal overlap may *increase* false memory susceptibility. Consecutive presentation of semantically associated study items may promote attention to the shared meaning between study items rather than specific, differentiating details. That is, temporal overlap may promote gist-level memory rather than verbatim memory (Brainerd & Reyna, 1990, 2002), which has been linked to false memory (Tun, Wingfield, Rosen, & Blanchard, 1998). Thus, temporal features may modulate false memory susceptibility in young and older adults.

Our hypothesis is that older adults rely more heavily on semantic versus episodic processing, which promotes false memory. To test our hypothesis, we conducted a behavioral study in which young (18 - 35) and older adult (60 - 85) participants studied word lists in anticipation of a memory test. We manipulated the degree of semantic overlap between words by varying the number of semantic associates, or words that share meaning, on the study list. We manipulated the degree of temporal overlap between semantically overlapping words by varying the number of intervening words between semantic associates. Following study, participants completed a recognition memory test composed of both study words and critical lures. We measured true and false memory rates as a function of age, temporal and semantic overlap. To the extent that older adults utilize semantic rather than temporal information to guide memory behavior, we expected to find that true and false memory rates differ as a function of temporal overlap strength only in young adults.

## Materials and Methods

### Transparency and Openness

The raw, deidentified data and the associated experimental and analysis codes used in this study will be available via the Long Term Memory lab website following publication (https://longtermmemorylab.com). The study design and analyses were preregistered (https://osf.io/tkhu9). We affirm that we report how we determined our sample size, any data exclusions, all manipulations, and all measures. Informed consent was obtained in accordance with the UVA Institutional Review Boards for Social and Behavioral Research and Health Sciences Research, and participants were compensated for their participation.

### Participants

Fifty young adult (28 female, age range = 18-35, mean age = 21.12 years) and fifty older adult (38 female, age range = 60-85, mean age = 70.38 years) fluent English speakers from the University of Virginia (UVA; Charlottesville, VA, USA) community participated. Young adult participants were recruited from flyers posted around the UVA campus and surrounding areas. Older adults were recruited through the Virginia Cognitive Aging Project, a longitudinal study of cognitive aging in adults across a wide age range (Salthouse, 2011). Our enrollment target was determined *a priori* based on young adult pilot data (N = 14) described in the preregistration report of this study (https://osf.io/tkhu9). All participants had normal or corrected-to-normal vision. All participants completed the Montreal Cognitive Assessment (MoCA); any participant who scored less than 26 was excluded, based on previous literature (Nasreddine et al., 2005). Data collection started and ended in 2024.

One young adult participant was excluded from the final dataset because they scored lower than 26 on the MoCA. Nine older adult participants were excluded from the final dataset: seven who scored lower than 26 on the MoCA, one who was younger than the lower age limit of 60 at the time of testing, and one who had poor task performance (false memory rate *>* 2.5 SDs of the mean of the full older adult dataset). Thus, data are reported for the remaining 49 young adult and 41 older adult participants. The raw, de-identified data and the associated experimental and analysis codes used in this study will be made available for access via the Long Term Memory laboratory website upon publication of this manuscript (https://longtermmemorylab.com).

### Recognition Task Experimental Design

Stimuli consisted of 1602 words, drawn from the Toronto Noun Pool (Friendly, Franklin, Hoffman, & Rubin, 1982). From this set, 386 words were randomly selected for each participant. Of these words, 288 were presented during the study phase. During the test phase, 108 study words were presented as targets and 108 unstudied words that semantically overlap with studied words (Roediger & McDermott, 1995) were presented as critical lures. 180 study words were not tested.

#### Study Phase

In each of 18 runs, participants viewed a list containing 16 words, yielding a total of 288 trials. During each trial, participants saw a single word presented for 2000 ms followed by a 1000 ms inter-stimulus interval (ISI; Figure 1). Participants were instructed to study the presented word in anticipation of a later memory test; participants did not make any behavioral responses. We manipulated the degree of temporal and semantic overlap between words on the study list. Semantic overlap was determined by the number of semantic associates each word had within a study list (Nelson, Zhang, & McKinney, 2001). Semantic associates were determined using word association space values (WAS; Nelson et al., 2001); semantic associates had a WAS value of 0.4 or greater (Long & Kahana, 2017). Half of the words overlapped semantically with three other words (strong semantic overlap; e.g. “actor,” “movie,” “screen,” “film”), one quarter overlapped semantically with only one other word (weak semantic overlap; e.g. “dog,” “cat), and one quarter did not overlap semantically with any other words (no semantic overlap; e.g. “virus”). Temporal overlap was determined based on the number of intervening words between semantic associates. Among the words with strong or weak semantic overlap, half of the words were presented consecutively with their semantic associates (strong temporal overlap; e.g. “table,” “chair”) and half were separated from their semantic associates by four words (weak temporal overlap; e.g. <u>“pail,”</u> “dog,” “shoe,” “fence,” “fan,” <u>“bucket”</u>). Each 16-item word list was constructed such that every list contained: one set of four semantic associates presented consecutively (strong semantic, strong temporal overlap), one set of four semantic associates presented with four intervening words between them (strong semantic, weak temporal overlap), one set of two semantic associates presented consecutively (weak semantic, strong temporal overlap), one set of two semantic associates presented with four intervening words between them (weak semantic, weak temporal overlap) and four unrelated words (no semantic overlap).

**Figure 1.**
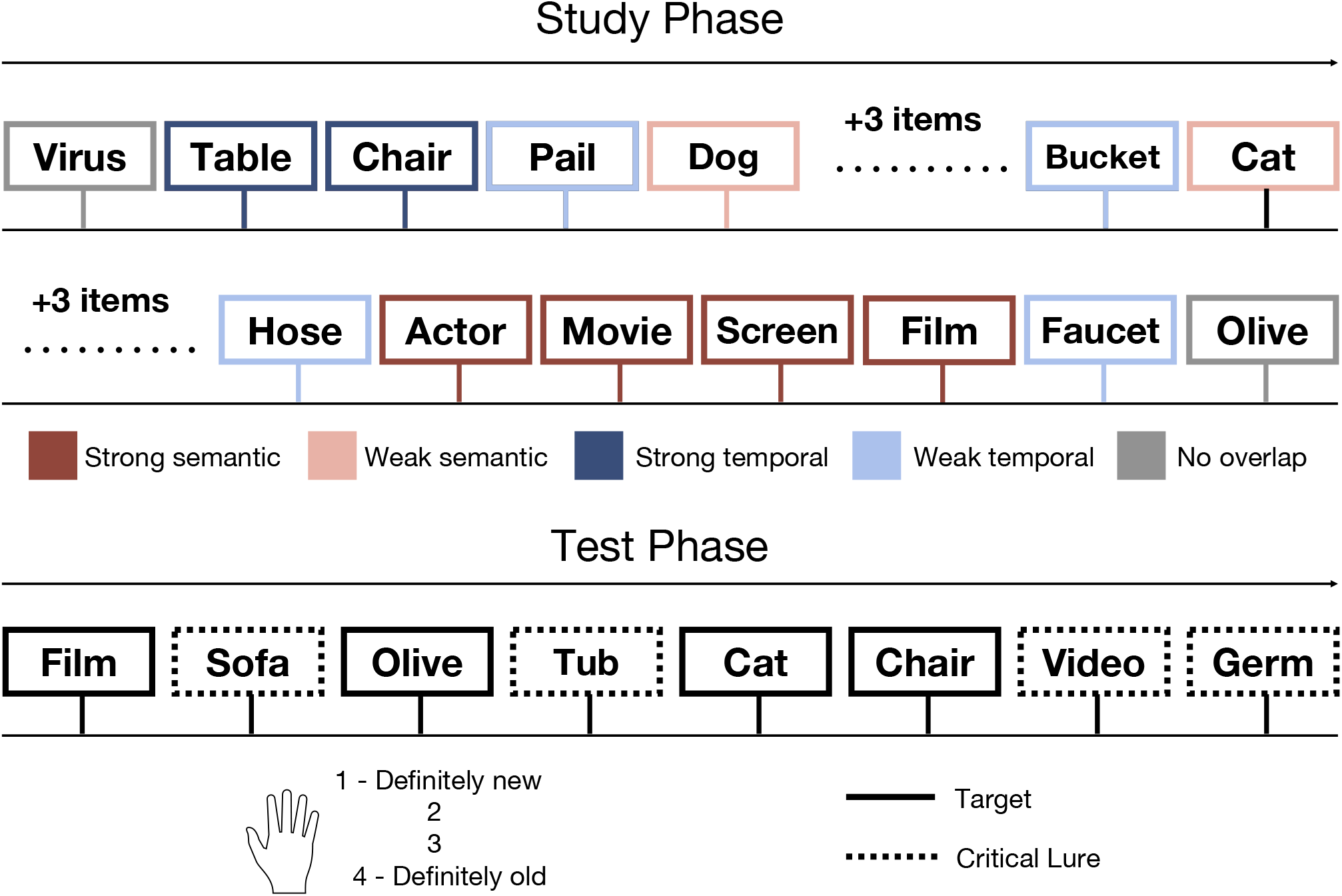
Experimental Design. During the study phase, participants studied individual words in anticipation of a later memory test and made no behavioral responses. Within each run, half of the words had strong semantic overlap (red; “actor,” “movie,” “screen,” “film”), one quarter had weak semantic overlap (pink; “dog,” “cat”) and one quarter had no semantic overlap (gray; “virus,” “olive”). Among those words with either strong or weak semantic overlap, half had strong temporal overlap (dark blue; “table,” “chair”) and half had weak temporal overlap (light blue; (weak temporal overlap; e.g. *“pail,”* “dog,” “shoe,” “fence,” “fan,” *“bucket”*). After 18 runs of 16-item word lists, participants completed a recognition test phase. On each test phase trial, participants saw either a target (solid line; a word that was presented during the study phase) or a critical lure (dashed line; a word that was not presented during the study phase but overlapped semantically with study words). The participants’ task was to make an old or new confidence judgment on a scale from one (definitely new) to four (definitely old). Lines and colors around the boxes are shown for illustrative purposes only and were not present during the actual experiment.

#### Test Phase

Following the 18 study runs, participants completed the recognition test phase. On each trial, participants viewed either a word which had been presented during the study phase (*target*) or an unstudied word that semantically overlapped with study words (*critical lure*; Roediger & McDermott, 1995). Participants’ task was to indicate their level of confidence that a given test probe was old or new on a scale from one (definitely new) to four (definitely old). Test trials were self-paced and responses could occur anytime after the stimulus onset. Participants received no feedback on the accuracy of their responses. Test trials were separated by a 1000 ms ISI. There were a total of 216 test trials, which consisted of 108 targets and 108 critical lures. In order to maintain an even number of targets and critical lures, only the last word of each “set” of semantic associates was presented as a target (e.g. if a participant studied “actor,” “movie,” “screen,” “film,” only “film” was presented during the test phase) and one semantically associated, unstudied word was presented as a critical lure (e.g. “video”). Test trial order was randomized.

### Behavioral Data Analyses

We assessed false memory rate by calculating the probability that a participant responded “old” (both high and low confidence) out of the total number of trials in which they were presented with a critical lure. We assessed hit rate by calculating the probability that a participant responded “old” (both high and low confidence) out of the total number of trials in which they were presented with a target. Hit and false alarm rates were calculated separately for our conditions of interest, namely: age group (young, older), semantic overlap strength (strong, weak) and temporal overlap strength (strong, weak). To evaluate false memory confidence, we separately calculated the probability that a participant responded “definitely old” or “unsure old” to calculate high and low confidence false memory rate, respectively. We also assessed memory discrimination (d′) between targets and lures. For each participant, we calculated d′ by subtracting the normalized false alarm rate (the percentage of critical lures that were incorrectly identified as “old”) from the normalized hit rate (the percentage of targets that were correctly identified as “old”) separately for items with each condition of interest and items with no semantic overlap. We calculated a d′ difference score for each participant by subtracting d′ for items with no semantic overlap from d′ values for each condition of interest.

### Statistical Analyses

#### Planned analyses

We used a series of mixed effects ANOVAs (meANOVAs) to evaluate the effects of age and temporal overlap strength on true and false memory. Specifically, we used separate meANOVAs to assess the effects of age group (young, older) and temporal overlap strength (strong, weak) on three dependent variables: (1) false memory rate for critical lures associated with strongly semantically overlapping study items, (2) high confidence false memory rate for critical lures associated with strongly semantically overlapping study items, and (3) hit rate for strongly semantically overlapping targets. For each of the three dependent variables, we used Bayes Factor analyses to compare models with and without the two-way interaction term of age and temporal overlap strength. We used repeated measures ANOVAs (rmANOVAs) to assess the effects of temporal and semantic overlap strength (strong, weak) on hit and false memory rates in older adults. We used post-hoc independent *t* -tests for age group comparisons and paired *t* -tests for within age condition comparisons.

#### Exploratory analyses

We conducted exploratory meANOVAs to assess the effects of age, temporal overlap strength, and semantic overlap strength on three dependent variables: (1) false memory rate, (2) hit rate and (3) d′ difference score. We conducted post-hoc paired *t* -tests to compare false memory rates as a function of semantic overlap strength across age. We conducted a post-hoc paired *t* -test to test the effect of semantic overlap strength on young adults’ hit rates for strongly temporally overlapping targets. We used Bayes Factor analyses to compare models of false memory rate and d′ difference score with and without the three-way interaction term of age, semantic and temporal overlap strength. We used an meANOVA to assess the effects of age, temporal overlap, semantic overlap, and confidence (high, low) on false memory rate. We used Bayes Factor analysis to compare models of false memory rate with and without the three-way interaction term of age, temporal overlap strength, and confidence. Separately for low and high confidence false memories, we used post-hoc paired *t* -tests to compare false memory rates as a function of temporal or semantic overlap strength.

## Results

### Over-reliance on semantic information increases false memory susceptibility

According to our hypothesis, older adults may rely more heavily on semantic versus episodic processing as compared to young adults. Thus, our first goal was to determine the extent to which young and older adults utilize temporal (e.g. episodic) information to reject critical lures. Following our pre-registration, we conducted a 2 *×* 2 mixed effects ANOVA (meANOVA) to evaluate the effects of age group (young, older) and temporal overlap strength (strong, weak) on false memory rate for critical lures associated with strongly semantically overlapping study items (Figure 2A). The ability to access shared temporal contextual information to reject critical lures should lead to lower false alarm rates for critical lures associated with strongly vs. weakly temporally overlapping study items. To the extent that older adults are less sensitive to temporal information in comparison to young adults, their false memory rates should not be modulated by temporal overlap strength. Therefore, we should find a two-way interaction between age group and temporal overlap strength. Alternatively, a lack of a two-way interaction between age group and temporal overlap strength would indicate that the two age groups are equally sensitive, or insensitive, to temporal information. We find a significant main effect of age group (*F* _1,88_ = 5.81, *p* = 0.018, 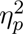), driven by a greater false memory rate for older (*M* = 0.30, *SD* = 0.17) compared to young (*M* = 0.22, *SD* = 0.11) adults. The main effect of temporal overlap strength was not significant (*F* _1,88_ = 0.04, *p* = 0.842, 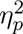), nor was the two-way interaction between age group and temporal overlap strength (*F* _1,88_ = 0.03, *p* = 0.856, 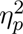). Bayes Factor analysis revealed that a model without the two-way interaction term (H_0_) is preferred to a model with the two-way interaction term (H_1_; BF_10_ = 0.21, moderate evidence for H_0_). These results indicate that under conditions of strong semantic overlap, false memory was modulated solely by age group, with no evidence for an influence of temporal overlap strength. That we do not find an interaction between age and temporal overlap strength provides evidence against our hypothesis and instead suggests that the use of temporal information to reject critical lures does not differ between age groups.

**Figure 2.**
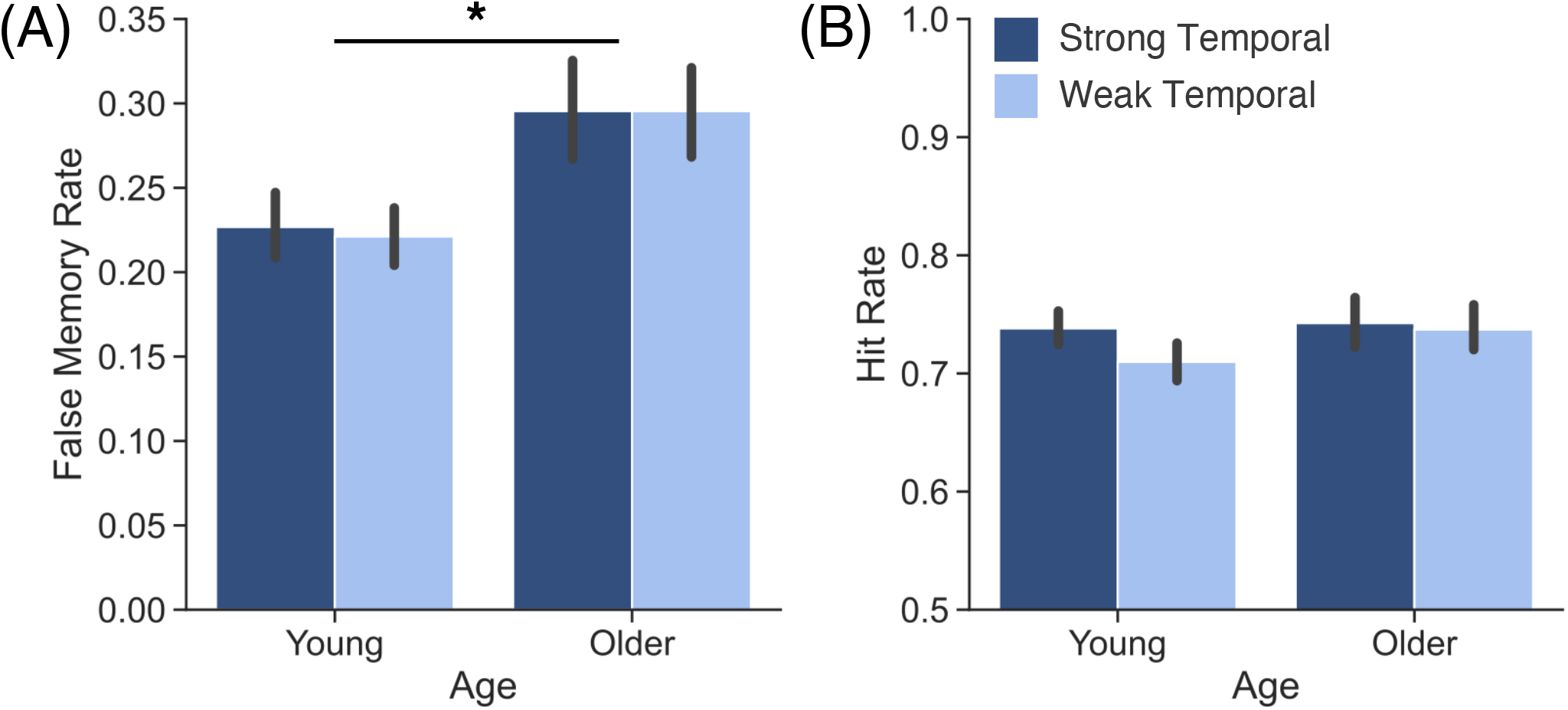
Influence of age and temporal overlap strength on memory behavior for words with strong semantic overlap. We assessed **(A)** false memory and **(B)** hit rate for words with strong semantic overlap as a function of age (young, older) and temporal overlap strength (strong, dark blue; weak, light blue). We find a significant main effect of age (*p* = 0.018) driven by greater false memory rates for older than young adults. Note that y-axes differ between panels A and B. \**p <* 0.05. Error bars denote standard error of the mean.

Given that young and older adults appear equally insensitive to temporal information when rejecting lures, we next sought to determine their relative sensitivity when endorsing targets. Following our preregistration, we conducted a 2 *×* 2 meANOVA to evaluate the effects of age group (young, older) and temporal overlap strength (strong, weak) on hit rate for strongly semantically overlapping targets (Figure 2B). The ability to access shared temporal contextual information to endorse targets should lead to greater hit rates for strongly versus weakly temporally overlapping targets. Thus, we should find a two-way interaction between age group and temporal overlap strength. The main effects of age group (*F* _1,88_ = 0.25, *p* = 0.617, 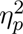) and temporal overlap strength were not significant (*F* _1,88_ = 1.25, *p* = 0.267, 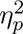). Consistent with our findings for false memory, we find that the two-way interaction between age group and temporal overlap strength on hit rate was not significant (*F* _1,88_ = 0.51, *p* = 0.478, 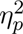). Bayes Factor analysis revealed that a model without the two-way interaction term (H_0_) is preferred to a model with the two-way interaction term (H_1_; BF_10_ = 0.18, moderate evidence for H_0_). Contrary to our hypothesis, we find no evidence for an interaction between the effects of age and temporal overlap strength on true memory, suggesting that young and older adults similarly utilize temporal contextual information to recognize targets.

Increased overlap across any dimension (semantic or temporal) may lead to less differentiated representations in older adults, regardless of their relative sensitivity to temporal versus semantic information. Relative to young adults, older adults’ neural representations tend to be de-differentiated or reduced in distinctiveness (Koen & Rugg, 2019; Folville et al., 2020; Hill, King, & Rugg, 2021; Pauley, Karlsson, & Sander, 2024; Srokova, Aktas, Koen, & Rugg, 2024). One consequence of de-differentiation is that current experiences can be more readily integrated with existing representations (Spreng & Schacter, 2012; Geerlings, Saliasi, Renken, Maurits, & Lorist, 2014; Amer, Anderson, Campbell, Hasher, & Grady, 2016; Rieck, Rodrigue, Boylan, & Kennedy, 2017), such that the representations of critical lures may become integrated with those of study items. As a result, we would expect to find that older adults have greater false memory rates for critical lures that are associated with strongly overlapping study items regardless of the dimension of said overlap. To test this possibility, we conducted a pre-registered 2 *×* 2 repeated measures ANOVA (rmANOVA) to evaluate the effects of temporal and semantic overlap strength on false memory rate in older adults only (Figure 3, Older Adults). If a general increase in overlap leads to de-differentiated representations, we would expect to find main effects of both semantic and temporal overlap, as well as an interaction between the two dimensions, whereby false memory rate is greatest for items that overlap across both dimensions. However, if our hypothesis is correct and older adults are specifically insensitive to temporal information, we would expect to find a main effect of semantic overlap with no main effect of temporal overlap and no interaction between temporal and semantic overlap. We find a significant main effect of semantic overlap strength (*F* _1,40_ = 11.21 *p* = 0.002, *η*^2^ = 0.22), driven by greater false memory rate for critical lures associated with strongly (*M* = 0.30, *SD* = 0.17) compared to weakly semantically overlapping study items (*M* = 0.25, *SD* = 0.18). The main effect of temporal overlap strength (*F* _1,40_ = 0.29 *p* = 0.593, *η*^2^ = 0.01) and the interaction between temporal and semantic overlap strength were not significant (*F* _1,40_ = 0.14 *p* = 0.708, *η*^2^ = 0.00). Bayes Factor analysis revealed that a model without the two-way interaction term (H_0_) is preferred to a model with the two-way interaction term (H_1_; BF_10_ = 0.22, moderate evidence for H_0_). These results suggest that older adults’ false memory susceptibility arises from an over-reliance on semantic versus episodic processing, rather than a more general de-differentiation of representations across dimensions.

**Figure 3.**
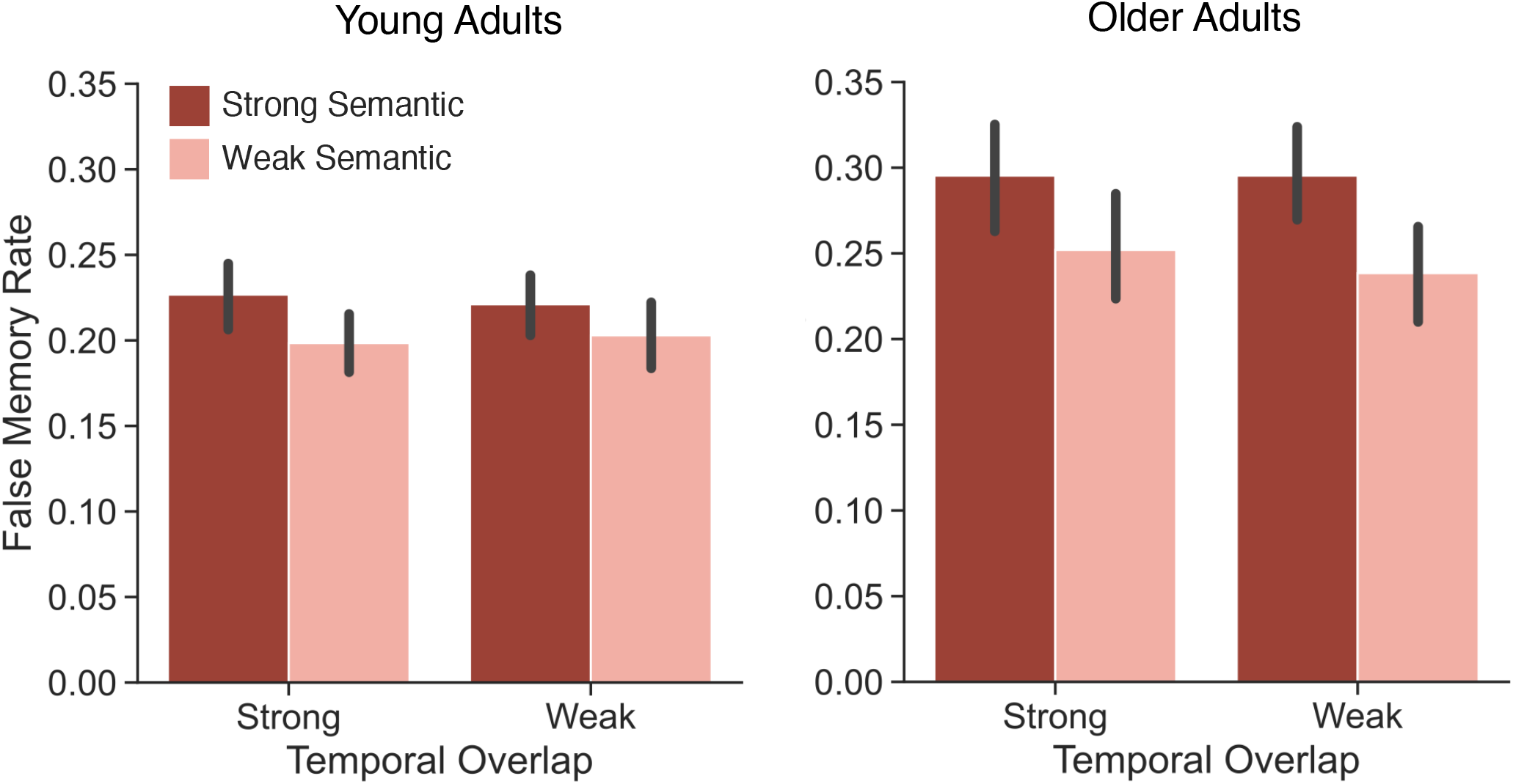
Influence of age, temporal overlap strength, and semantic overlap strength on false memory. We assessed false memory rate as a function of age (young, older), temporal overlap strength (strong, weak) and semantic overlap strength (strong, red; weak, pink). We find a significant main effect of semantic overlap strength (*p <* 0.001) driven by greater false memory rate for words with strong compared to weak semantic overlap. Error bars denote standard error of the mean.

Our findings indicate that young and older adults may have similar access to temporal information to endorse targets and reject critical lures when there is an abundance of semantic overlap between study items. That is, we specifically examined age-related effects of temporal overlap strength on strongly *semantically* overlapping items. The rationale for this approach is that prior work has shown that both young and older adults are highly susceptible to false memories when many study items semantically overlap (Koutstaal & Schacter, 1997). Thus, insofar as participants utilize temporal information to guide memory behavior, manipulating the temporal overlap strength of strongly semantically overlapping items may impact false alarm rates. However, the presence of many semantically associated study items may increase the relative salience of semantic information to the point of biasing both young and older adults to selectively attend to semantic, rather than temporal, information. Explicit task instructions to perform a semantic judgment and/or the presence of semantic associates can selectively focus attention to semantic information, often at the expense of temporal information (Long & Kahana, 2017; Moore & Long, 2024). Therefore, we next examined the influence of temporal overlap strength on both strongly and weakly semantically overlapping study items.

We conducted an exploratory analysis to determine the extent to which strong versus weak overlap in either dimension (temporal or semantic) modulates false memory in young and older adults. To the extent that older adults rely more heavily on semantic versus episodic processing, we should find greater false memory rates for critical lures associated with strongly vs. weakly semantically overlapping study items, with no influence of temporal overlap strength. In contrast, young adults may rely on temporal information, especially when semantic contextual overlap is weak or absent. Thus, we should find a three-way interaction between age, semantic overlap strength, and temporal overlap strength. Alternatively, the two age groups may be equally biased toward semantic and away from episodic information such that both older and young adults are more susceptible to false memories for critical lures associated with strongly semantically overlapping study items, regardless of temporal overlap strength.

We conducted an exploratory 2 *×* 2 *×* 2 meANOVA to evaluate the effects of age group (young, older), temporal overlap strength (strong, weak) and semantic overlap strength (strong, weak) on false memory rate (Figure 3). A three-way interaction would indicate that young and older adults differentially utilize temporal and semantic information when endorsing critical lures. Alternatively, a significant main effect of semantic overlap strength with no three-way interaction would indicate that false memory rate is modulated by semantic overlap strength similarly for both age groups, regardless of temporal overlap. We report the full ANOVA results in Table 1 and highlight key findings here. The three-way interaction between age, temporal overlap strength and semantic overlap strength was not significant (BF_10_ = 0.26, moderate evidence for H_0_). We find a significant main effect of semantic overlap strength, driven by greater false memory rate for critical lures associated with strongly (*M* = 0.26, *SD* = 0.14) compared to weakly (*M* = 0.22, *SD* = 0.15) semantically overlapping study items. We find no evidence that temporal overlap influences false memory in young or older adults, contrary to our hypothesis. These results instead indicate that both age groups are sensitive to semantic overlap strength such that false memories are greater for critical lures associated with strongly compared to weakly semantically overlapping study words. Our findings suggest that when events overlap temporally and semantically, young and older adults rely predominantly on semantic information when endorsing, or rejecting, critical lures.

**Table 1.** Analysis of variance for the effects of age group, temporal overlap strength, and semantic overlap strength on false memory rate.

| Effect | $F_{1,88}$ | $p$ | $\eta_p^2$ |
| --- | --- | --- | --- |
| Main effect of age group | 4.09 | **0.046** | 0.04 |
| Main effect of temporal overlap | 0.13 | 0.719 | 0.00 |
| Main effect of semantic overlap | 14.98 | **<0.001** | 0.15 |
| Interaction of age group × temporal overlap | 0.11 | 0.743 | 0.00 |
| Interaction of age group × semantic overlap | 2.13 | 0.145 | 0.02 |
| Interaction of temporal overlap × semantic overlap | 0.00 | 0.979 | 0.00 |
| Interaction of age group × temporal overlap × semantic overlap | 0.26 | 0.614 | 0.00 |

*Bold values indicate $p < 0.05$ .*

### False memory confidence scales with overlap strength

We did not find age-related differences in false memory rate as a function of overlap strength; however, our inability to detect such differences may be the result of collapsing over confidence. That is, for all previous analyses, we included both low confidence (“unsure old” responses) and high confidence (“definitely old” responses) false alarms, which may obscure underlying age-related differences between false memories for critical lures associated with strongly vs. weakly temporally overlapping study items. Following our pre-registration, we performed a 2 *×* 2 meANOVA to evaluate the effects of age group (young, older) and temporal overlap strength (strong, weak) on high confidence false memory rates for critical lures associated with strongly semantically overlapping study items (Table 2; Figure 4A, High Confidence). If young and older adults differ in their use of temporal information, we should find a two-way interaction. Specifically, we would expect to find that young adults more confidently reject critical lures associated with strongly temporally overlapping study items. In contrast, we would expect to find that older adults’ high confidence false memory rates are not modulated by temporal overlap strength. Alternatively, a main effect of temporal overlap strength, and no two-way interaction with age, would indicate that temporal overlap strength modulates behavior similarly for both young and older adults. We find a significant main effect of temporal overlap strength, driven by a greater high confidence false memory rate for critical lures associated with strongly (*M* = 0.16, *SD* = 0.15) compared to weakly (*M* = 0.13, *SD* = 0.13) temporally overlapping study items. The interaction between age and temporal overlap strength was not significant. Bayes Factor analysis revealed that a model without the two-way interaction term (H_0_) is preferred to a model with the three-way interaction term (H_1_; BF_10_ = 0.20, moderate evidence for H_0_). These results suggest that for both young and older adults, consecutively encountering multiple study items that overlap semantically may promote attention to the semantic dimension, leading to higher confidence false memories.

**Table 2.** Analysis of variance for the effects of age group and temporal overlap strength on high confidence false memory rate for words with strong semantic overlap.

| Effect | $F_{1,88}$ | $p$ | $\eta_p^2$ |
| --- | --- | --- | --- |
| Main effect of age | 13.29 | **<0.001** | 0.13 |
| Main effect of temporal overlap | 5.29 | **0.024** | 0.06 |
| Interaction of age × temporal overlap | 0.02 | 0.901 | 0.00 |

*Bold values indicate $p < 0.05$ .*

**Figure 4.**
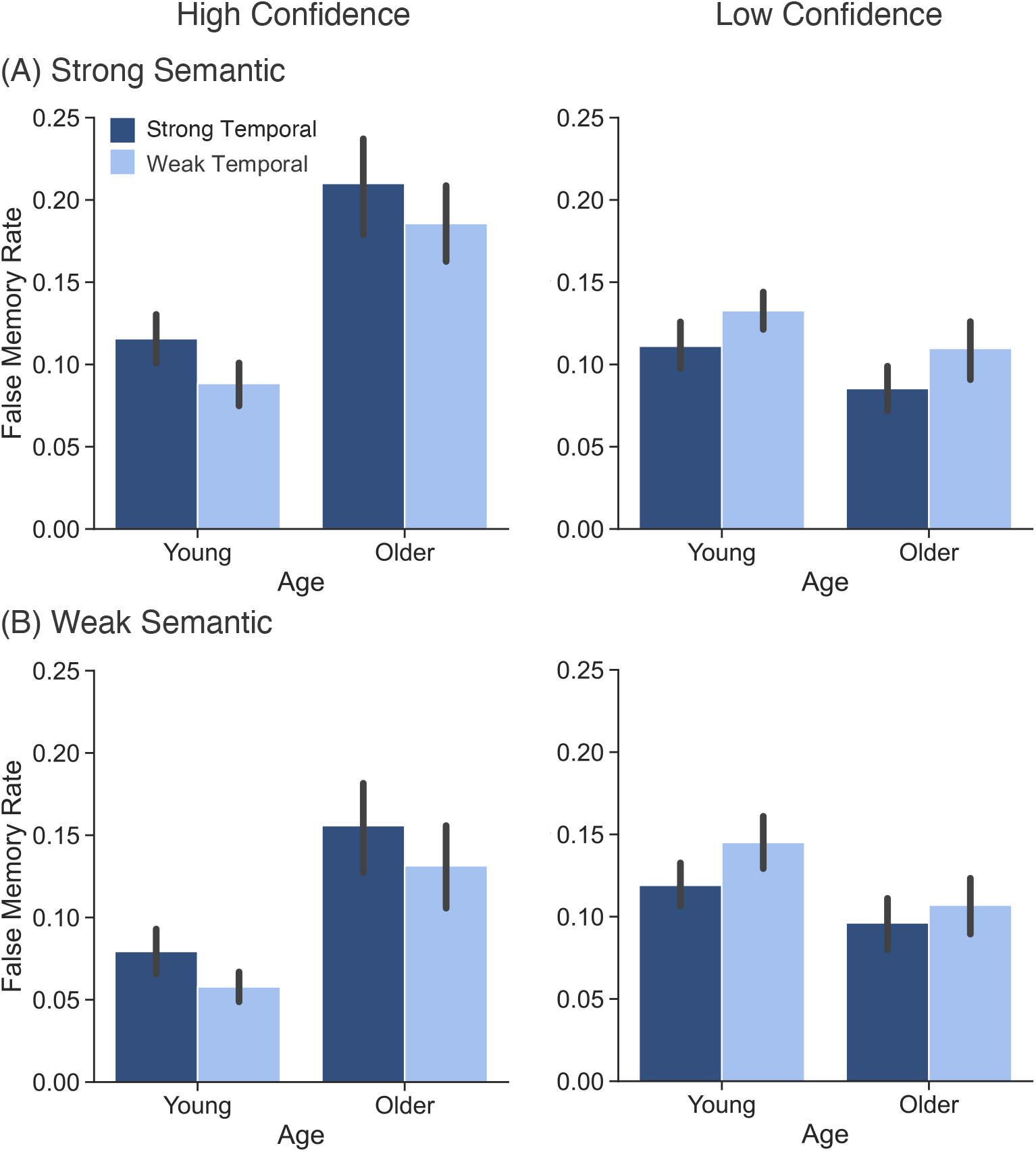
Influence of age, temporal overlap strength, semantic overlap strength, and confidence on false memory. We assessed false memory rate as a function of age (young, older), temporal overlap strength (strong, dark blue; weak, light blue) and confidence (high, low) for critical lures associated with **(A)** strongly and **(B)** weakly semantically overlapping study items. Error bars denote standard error of the mean.

The results of the prior analysis suggest that young and older adults do not differentially utilize temporal information when they confidently endorse critical lures. However, given evidence that older adults typically have higher confidence in their false memories than young adults (Dodson & Krueger, 2006; Shing, Werkle-Bergner, Li, & Lindenberger, 2009), separately considering low and high confidence false memories may be more appropriate for capturing age-related differences in reliance on temporal versus semantic information. Thus, we performed an exploratory analysis in which we used a 2 *×* 2 *×* 2 *×* 2 meANOVA to evaluate the effects of age group (young, older), temporal overlap strength (strong, weak), semantic overlap strength (strong, weak) and confidence (high, low) on false memory rate for critical lures (Figure 4). If young adults are more sensitive to temporal information than older adults, we would expect to find that young adults have more high confidence false memories, and fewer low confidence false memories, for critical lures associated with strongly compared to weakly temporally overlapping study items. In contrast, older adults’ high and low confidence false memory rates should not be modulated by temporal overlap strength. In this case, a three-way interaction between age, temporal overlap strength and confidence would suggest that young and older adults use temporal information differently when presented with critical lures. Alternatively, if false memory confidence scales with the degree of overlap between study items regardless of dimension for both age groups, we would expect to find two-way interactions between temporal strength and confidence and semantic strength and confidence with no three-way interactions between temporal strength, confidence and age or semantic strength, confidence and age. We report the full ANOVA results in Table 3 and highlight key findings here.

**Table 3.** Analysis of variance for the effects of age group, temporal overlap strength, semantic overlap strength, and confidence on false memory rate.

| Effect | $F_{1,88}$ | $p$ | $\eta_p^2$ |
| --- | --- | --- | --- |
| Main effect of age group | 4.09 | **0.046** | 0.04 |
| Main effect of temporal overlap | 0.13 | 0.719 | 0.00 |
| Main effect of semantic overlap | 14.98 | **<0.001** | 0.15 |
| Main effect of confidence | 0.38 | 0.535 | 0.00 |
| Interaction of age group × temporal overlap | 0.11 | 0.743 | 0.00 |
| Interaction of age group × semantic overlap | 2.13 | 0.148 | 0.02 |
| Interaction of age group × confidence | 12.96 | **0.001** | 0.13 |
| Interaction of temporal overlap × confidence | 15.07 | **<0.001** | 0.15 |
| Interaction of semantic overlap × confidence | 17.87 | **<0.001** | 0.17 |
| Interaction of temporal overlap × semantic overlap | 0.00 | 0.979 | 0.00 |
| Interaction of age group × temporal overlap × confidence | 0.07 | 0.793 | 0.00 |
| Interaction of age group × semantic overlap × confidence | 0.37 | 0.542 | 0.00 |
| Interaction of age group × semantic overlap × temporal overlap | 0.26 | 0.614 | 0.00 |
| Interaction of age group × semantic × temporal × confidence | 0.08 | 0.782 | 0.00 |

*Bold values indicate $p < 0.05$ .*

We find a significant two-way interaction between temporal strength and confidence driven by greater high confidence false memories for critical lures associated with strongly (*M* = 0.14, *SD* = 0.14) compared to weakly (*M* = 0.11, *SD* = 0.12) temporally overlapping study items (*t* _89_ = 3.70, *p <* 0.001, *d* = 0.19) and greater low confidence false memories for critical lures associated with weakly (*M* = 0.13, *SD* = 0.09) compared to strongly (*M* = 0.10, *SD* = 0.08) temporally overlapping study items (*t* _89_ = 2.55, *p* = 0.013, *d* = 0.24). Thus, both young and older adults are more susceptible to high confidence false memories when associated study items are presented consecutively. Both age groups are more likely to demonstrate low confidence in their false memories when study items are not presented consecutively. We find a significant two-way interaction between semantic strength and confidence driven by greater high confidence false memories for critical lures associated with strongly (*M* = 0.15, *SD* = 0.13) compared to weakly (*M* = 0.10, *SD* = 0.13) semantically overlapping study items (*t* _89_ = 5.42, *p <* 0.001, *d* = 0.33). Low confidence false memory rate did not significantly differ between critical lures associated with strongly (*M* = 0.11, *SD* = 0.09) or weakly (*M* = 0.12, *SD* = 0.09) semantically overlapping study items (*t* _89_ = 1.05, *p* = 0.299, *d* = 0.09). Neither three-way interactions were significant and Bayes Factor analyses yielded moderate evidence in support of models without the three-way interaction terms (temporal *×* age *×* confidence, BF_10_ = 0.18; semantic *×* age *×* confidence, BF_10_ = 0.19). These results indicate that both young and older adults have higher confidence in their false memories when study items overlap, regardless of overlap dimension.

### Young and older adults differentially utilize temporal information to support true memory

Having failed to detect age differences in the effects of episodic information on false memory, we next sought to determine the extent to which young and older adults utilize episodic versus semantic information to support true memory. The degree to which individuals attend to semantic information should have a differential impact on recognition given that not all items have strong semantic overlap and attending to semantic information can negatively impact later memory when a study item has few or no semantic associates on the study list (Long & Kahana, 2017). Furthermore, we expect that the “default” orientation of attention differs across age. Namely, young adults’ relatively preserved episodic memory should lead to a “default” tendency to attend to the episodic or event-defining, distinct features of events (Healey, 2018), whereas older adults’ diminished episodic memory (Wingfield & Kahana, 2002) and corresponding semantic bias (Ofen & Shing, 2013) should shift their “default” tendency to instead attend to the semantic dimension of events. These combined effects – the utility of semantic information and the tendency to attend to it – should produce differential hit rates across age and condition. To the extent that older adults rely more heavily on semantic versus episodic processing than young adults, we would expect to find a three-way interaction between age group, temporal overlap strength and semantic overlap strength on hit rate.

We conducted an exploratory 2 *×* 2 *×* 2 meANOVA to evaluate the effects of age group, temporal overlap strength and semantic overlap strength on hit rate (Figure 5). We report the full ANOVA results in Table 4 and highlight key findings here. We find a marginal three-way interaction between age group, temporal overlap strength, and semantic overlap strength, providing support for our hypothesis. Specifically, young adults have higher hit rates for strongly (*M* = 0.74, *SD* = 0.14) compared to weakly (*M* = 0.67, *SD* = 0.16) semantically overlapping targets that are *strongly* temporally overlapping (*t* _48_ = 3.41, *p* = 0.001, *d* = 0.46). Young adult hit rate does not significantly differ between strongly (*M* = 0.71, *SD* = 0.16) compared to weakly (*M* = 0.68, *SD* = 0.16) semantically overlapping targets that are *weakly* temporally overlapping (*t* _48_ = 1.50, *p* = 0.140, *d* = 0.17). This result means that young adults are more likely to recognize “film” than “chair” but equally likely to recognize “faucet” and “cat.” Older adults have a higher hit rate for strongly (*M* = 0.74, *SD* = 0.17) compared to weakly (*M* = 0.65, *SD* = 0.21) semantically overlapping targets that are *weakly* temporally overlapping (*t* _40_ = 4.92, *p <* 0.001, *d* = 0.45). Older adult hit rate does not significantly differ between strongly (*M* = 0.74, *SD* = 0.19) compared to weakly (*M* = 0.70, *SD* = 0.18) semantically overlapping targets that are *strongly* temporally overlapping (*t* _40_ = 1.68, *p* = 0.101, *d* = 0.22). This result means that older adults are more likely to recognize “faucet” than “cat” but equally likely to recognize “film” and “chair.” Thus, young and older adults appear to differentially leverage episodic information to support true memory. Young adults benefit from items overlapping both semantically and episodically, which may work to orient their attention to the semantic dimension. In contrast, older adults benefit from semantic overlap specifically when episodic information is weak.

**Figure 5.**
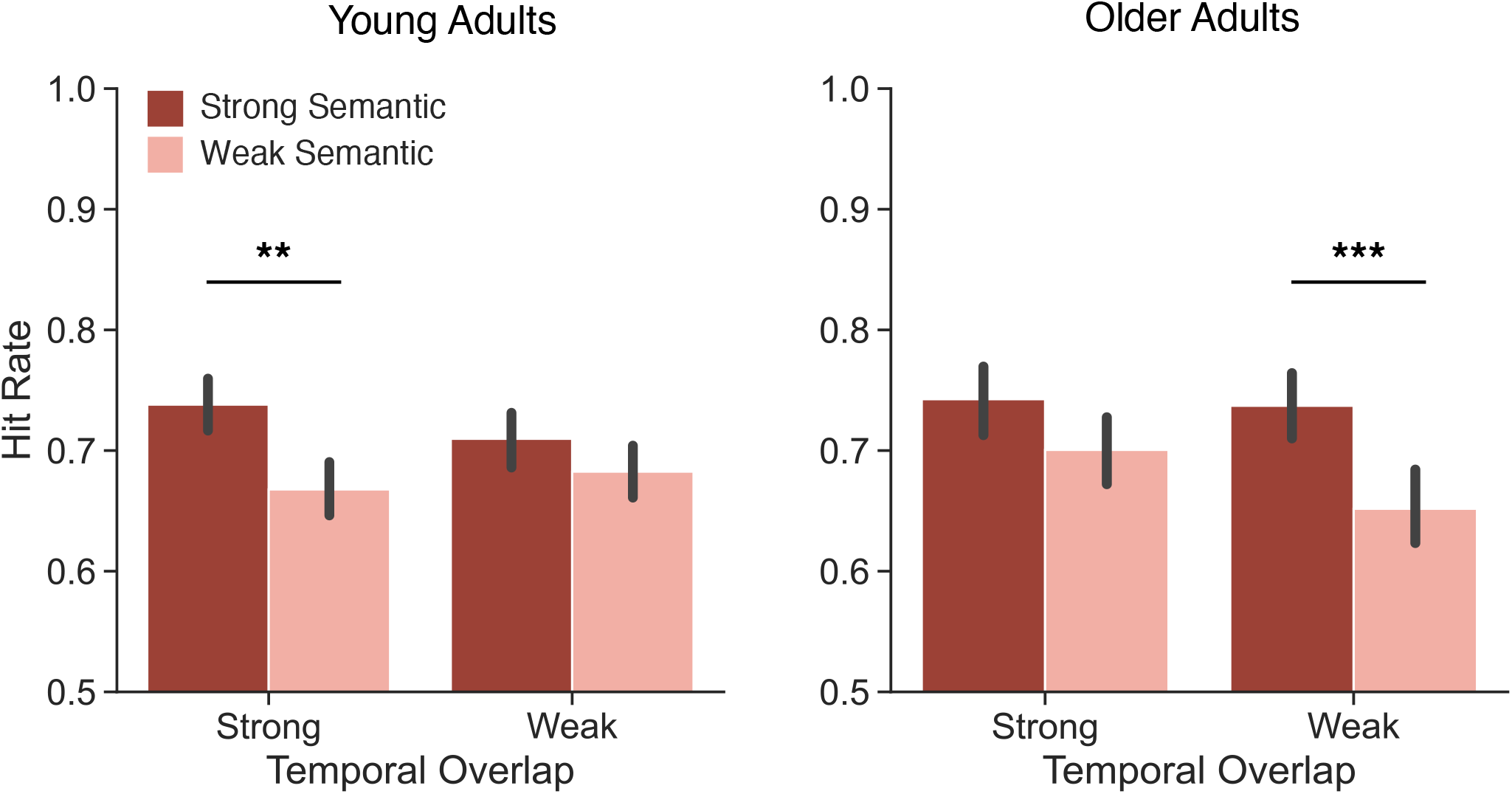
Influence of age and temporal and semantic overlap strength on true memory. We assessed hit rate as a function of age (young, older), temporal overlap strength (strong, weak) and semantic overlap strength (red represents strong semantic overlap; pink represents weak semantic overlap). We find a marginal interaction between age group, temporal overlap strength, and semantic overlap strength (*p* = 0.055) whereby older adults have a diminished hit rate for weak compared to strong semantic overlap for words with weak temporal overlap (*p <* 0.001) and young adults have a diminished hit rate for weak compared to strong semantic overlap for items with strong temporal overlap (*p* = 0.001). \*\**p <* 0.01, \*\*\**p <* 0.001. Error bars denote standard error of the mean.

**Table 4.** Analysis of variance for the effects of age group, temporal overlap strength, and semantic overlap strength on hit rate.

| Effect | $F_{1,88}$ | $p$ | $\eta_p^2$ |
| --- | --- | --- | --- |
| Main effect of age group | 0.07 | 0.790 | 0.00 |
| Main effect of temporal overlap | 2.02 | 0.159 | 0.02 |
| Main effect of semantic overlap | 36.18 | **<0.001** | 0.29 |
| Interaction of age group × temporal overlap | 0.80 | 0.373 | 0.01 |
| Interaction of age group × semantic overlap | 0.65 | 0.423 | 0.01 |
| Interaction of temporal overlap × semantic overlap | 0.03 | 0.868 | 0.00 |
| Interaction of age group × temporal overlap × semantic overlap | 3.78 | **0.055** | 0.04 |

*Bold values indicate $p < 0.05$ .*

Our results indicate that true memory differs by age as a function of temporal and semantic overlap strength. However, the previous analysis does not account for the possibility that there may be age-related differences in memory discriminability. We expect that the age-related differences we observe arise from older adults attending more to semantic than episodic features relative to young adults. However, given evidence that older adults respond more liberally on recognition memory tasks (Fraundorf, Hourihan, Peters, & Benjamin, 2019), older adults may be biased to respond “old” to all test words regardless of probe type (target or critical lure). To adjudicate between these possibilities, we assessed memory discrimination as a function of age. Specifically, we calculated a d′ difference score by subtracting d′ for items with no semantic overlap (and also, therefore, no temporal overlap) from d′ for each condition of interest. We conducted an exploratory 2 *×* 2 *×* 2 meANOVA to evaluate the effects of age group (young, older), temporal overlap strength (strong, weak) and semantic overlap strength (strong, weak) on d′ difference score (Figure 6). We find no significant main effects or interactions (all *p*s *>* 0.180; three-way interaction BF_10_ = 0.31, moderate evidence for H_0_; Table 5), indicating that memory discriminability is consistent across age groups and temporal/semantic overlap conditions. These findings suggest that the age-related differences in true memory are the result of differences in the relative reliance on semantic versus episodic processing, rather than response biases. Our interpretation is that young and older adults differentially attend to semantic versus episodic information to support true memory.

**Figure 6.**
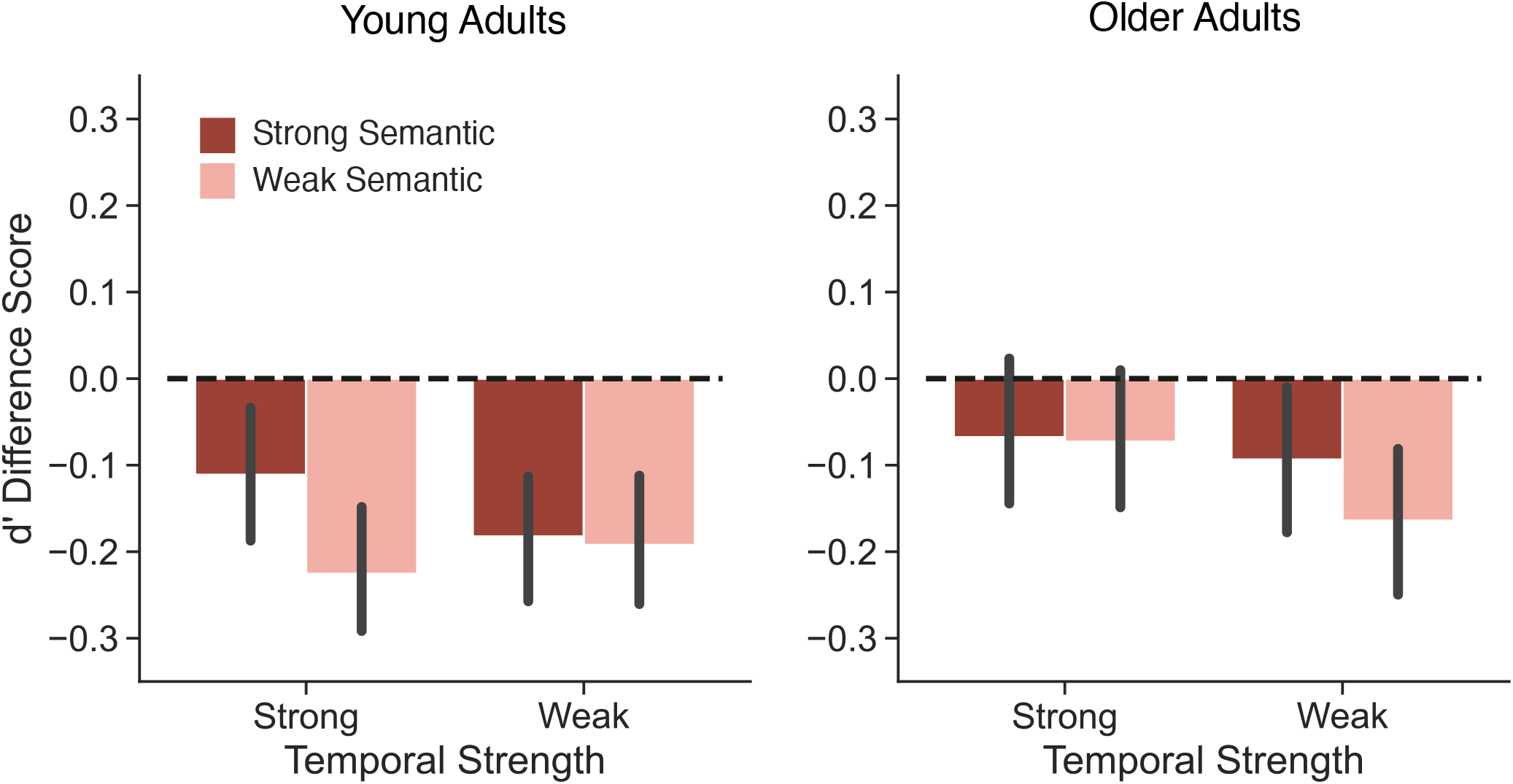
Influence of age and temporal and semantic overlap strength on memory discrimination. We calculated d′ difference score by subtracting d′ for items with no semantic overlap from d′ for each condition of interest. We then assessed d′ difference score as a function of age (young, older), temporal overlap strength (strong, weak) and semantic overlap strength (red represents strong semantic overlap; pink represents weak semantic overlap). We find no significant main effects or interactions (all *p*s *>* 0.180). Error bars denote standard error of the mean.

**Table 5.** Analysis of variance for the effects of age group, temporal overlap strength, and semantic overlap strength on d′ difference score.

| Effect | $F_{1,88}$ | $p$ | $\eta_p^2$ |
| --- | --- | --- | --- |
| Main effect of age group | 0.96 | 0.330 | 0.01 |
| Main effect of temporal overlap | 0.53 | 0.470 | 0.01 |
| Main effect of semantic overlap | 1.82 | 0.180 | 0.02 |
| Interaction of age group × temporal overlap | 0.15 | 0.700 | 0.00 |
| Interaction of age group × semantic overlap | 0.10 | 0.751 | 0.00 |
| Interaction of temporal overlap × semantic overlap | 0.08 | 0.784 | 0.00 |
| Interaction of age group × temporal overlap × semantic overlap | 0.73 | 0.394 | 0.01 |

*Bold values indicate $p < 0.05$ .*

## Discussion

The goal of the current study was to measure the extent to which differential reliance on semantic compared to episodic processing contribute to increased false memory in older adults. We directly tested the hypothesis that older adults rely more heavily on semantic versus episodic processing, which promotes false memory. We collected behavioral data in young and older adults performing an old/new recognition memory task in which we manipulated the degree of semantic and temporal overlap between study words and measured true and false memory rates. We report three key findings. First, we find that reliance on semantic information, rather than episodic information, increases false memory rates for both young and older adults. Second, we show that young and older adults have more high confidence false memories when there is strong overlap between study items, regardless of overlap dimension. Finally, we find that young and older adults differentially utilize episodic information to support true memory. Taken together, these results suggest that young and older adults may differentially rely on semantic versus episodic information to support memory behavior.

We find that both young and older adults are more susceptible to false memory when stimuli overlap semantically with no apparent influence of temporal overlap on false memories. Given evidence that older adults tend to rely on gist-level, conceptual information as opposed to specific, distinguishing details (Koutstaal & Schacter, 1997; Koutstaal, Schacter, Galluccio, & Stofer, 1999), our finding that older adults’ false memory rates appear unaffected by temporal overlap was in line with our expectations. However, whereas we predicted that temporal overlap strength would influence false memory rate in young adults, we instead found that like older adults, false memory rates in young adults were not affected by temporal overlap. The lack of age-related effects of temporal overlap strength on false memory rates may be the result of a trade-off between attending to the semantic and episodic dimensions of an event. Our prior work shows that attending to semantic information comes at the expense of attending to episodic information (Long & Kahana, 2017). Thus, it is possible that any degree of semantic overlap orients attention toward the semantic dimension of study items (Moore & Long, 2024). Similarly, ‘weak’ temporal overlap as defined in the current paradigm may have been insufficiently distant in time to elicit temporal overlap strength effects selectively in young adults. Prior work suggests that the temporal information associated with a given item becomes more distinct from that of other items at longer timescales (Bower, 1972; Estes, 1955; Melton, 1970) and across study lists (Unsworth, Brewer, & Spillers, 2013; Wahlheim & Huff, 2015). Thus, operationalizing ‘weak’ temporal overlap as a separation of only four intervening items within a list may have impeded our ability to detect age-related temporal effects. Broadly, our findings are consistent with prior work demonstrating that young adults are more likely to falsely remember critical lures when study items semantically overlap (Gutchess & Schacter, 2012) and that young adults’ likelihood of experiencing false memory increases as more semantically overlapping items are presented during study (Koutstaal & Schacter, 1997).

In both young and older adults, increased overlap between events leads to more high confidence false memories. Specifically, we show that both age groups have more high confidence false memories for critical lures associated with strongly compared to weakly overlapping study items, regardless of whether those items overlap semantically or temporally. The tendency to provide high confidence “old” responses to strongly overlapping critical lures may be due to attentional shifts between gist vs. verbatim information, a build up of proactive interference, and/or alterations in metamemory abilities.

According to Fuzzy Trace Theory, false memory occurs when study items’ gist representations are stronger than their corresponding verbatim representations, enabling endorsement of critical lures that share gist (Tun et al., 1998). This interpretation is supported by evidence demonstrating that in young adults, false memories can be suppressed when participants are directed to focus on the distinctive details – verbatim representations – of individual items (Schacter, Israel, & Racine, 1999; Gallo, Roberts, & Seamon, 1997). In the current study, the high degree of overlap between study items, as well as the absence of instructions directing participants to attend to distinguishing details of study items, may have lead young adults to form more ‘older adult-like,’ gist-based representations. Strongly overlapping study items may promote attention to the gist-level information shared between study items rather than the verbatim-level details that enable differentiation between items (Brainerd & Reyna, 1990, 2002). Thus, when semantic associates appear multiple times throughout the study list (strong semantic overlap) and/or occur in quick succession (strong temporal overlap), gist representations may be reinforced such that participants become more susceptible to high confidence false memories.

Alternatively, overlap between study items may promote a build up of proactive interference, influencing false memory confidence. Proactive interference is a phenomenon in which past information disrupts the processing of new information (Szpunar, McDermott, & Roediger, 2008). As a result, performance decreases as more overlapping items are encountered (Underwood, 1957; Postman & Keppel, 1977; Szpunar et al., 2008). Prior work shows that when new study items overlap with old study items, proactive interference makes it more difficult to distinguish between items, increasing false memory susceptibility (Jacoby, Wahlheim, Rhodes, Daniels, & Rogers, 2010). Thus, if proactive interference builds up – either as a result of the presentation of multiple semantic associates, the close temporal proximity of semantic associates, or both – the resulting difficulty in distinguishing between items may lead to higher confidence false memories.

Finally, metamemory, or awareness of one’s own memory (Schneider & Laurion, 1993), may have been impacted by our study design. Whereas young adults’ metamemory abilities tend to be well-calibrated – young adults tend to provide higher confidence ratings for correct than incorrect responses (e.g. Roebers, 2002) – this calibration between confidence and accuracy generally declines with increasing task difficulty (Lichtenstein & Fischhoff, 1977; Pressley & Ghatala, 1988; Schneider & Laurion, 1993). Insofar as increased overlap between study items constitutes increased task difficulty, young and older adults alike may be less able to accurately calibrate confidence and accuracy, leading to more high confidence false memories.

We find age differences in the impact of semantic vs. temporal overlap on true memory. Specifically, we show that young adults have higher hit rates when a series of four semantic associates occur consecutively, compared to when only two semantic associates occur consecutively. In contrast, older adults have higher hit rates when a series of four semantic associates occur non-consecutively, compared to when two semantic associates occur non-consecutively. These findings are in line with our expectation that aging shifts the relative reliance on episodic versus semantic memory. Young adults leverage temporal information to support memory (Kahana, 1996). In free recall tasks, young adults are more likely than older adults to consecutively recall words from neighboring positions on the study list – reflecting attention toward temporal information (Golomb, Peelle, Addis, Kahana, & Wingfield, 2008). This evidence is consistent with our expectation that young adults have a “default” tendency to attend to the unique, episodic or spatiotemporal features that distinguish one event from another (Healey, 2018). In the current task, however, the presence of many semantic associates may work to ‘push’ young adults to attend more to the semantic dimension, making them older-adult like with regards to false alarms. At the same time, the presence of many semantic associates alone may not be sufficient to push young adults to focus entirely on semantic information, as we infer that they are able to further orient to the semantic dimension when presented with four consecutive semantic associates given the increased hit rates for these items. Thus, young adults may attend to episodic information unless task conditions promote attention to semantic information.

In contrast to young adults’ presumed tendency to focus on episodic features/information, older adults have diminished episodic memory (Wingfield & Kahana, 2002) and a bias toward semantic memory (Ofen & Shing, 2013). These age-related memory changes may reflect a shift in older adults’ “default” orientation such that they preferentially attend to the semantic dimension of events. Older adults have difficulty inhibiting retrieval of semantic information from memory (Wynn, Ryan, & Moscovitch, 2020) and rely on semantic rather than episodic memory, even when doing so is contrary to task goals (Lalla, Tarder-Stoll, Hasher, & Duncan, 2022). However, older adults’ orientation of attention toward the semantic dimension may often be adaptive. Semantic processing generally promotes true memory; when participants study semantically overlapping items, memory is typically improved at test (Jacoby & Wahlheim, 2013; Tullis, Benjamin, & Ross, 2014; McKinley, Ross, & Benjamin, 2019; McKinley & Benjamin, 2020; Antony et al., 2022; Moore & Long, 2024). Older adults may thus rely on semantic processing to compensate for diminished episodic fidelity. This compensatory account is consistent with neuroimaging evidence that has revealed age-related changes in the default mode network (DMN; Raichle et al., 2001). The DMN supports semantic processing (Binder, Desai, Graves, & Conant, 2009) and becomes increasingly coupled with prefrontal control regions in aging (Turner & Spreng, 2015; Spreng & Turner, 2019), enabling older adults to use semantic information to maintain task performance in the face of age-related cognitive declines (Adnan, Beaty, Lam, Spreng, & Turner, 2019; Spreng & Turner, 2019; Spreng et al., 2014). As a result of increased DMN-prefrontal control region coupling, neural representations of stimuli become less distinct – de-differentiated – in older adults (Koen & Rugg, 2019; Folville et al., 2020; Hill et al., 2021; Pauley et al., 2024; Srokova et al., 2024) such that semantically overlapping stimuli can be integrated with existing representations (Spreng & Schacter, 2012; Geerlings et al., 2014; Amer et al., 2016; Rieck et al., 2017). Thus, a shift in default attention to semantic information may reflect a compensatory mechanism recruited by older adults to maintain cognitive performance amidst episodic memory impairments.

The age-related dissociation in the use of episodic and semantic information to support true memory could be explained by differential processing during either study and/or test. Semantic overlap can modulate the orientation of attention – and the neural mechanisms recruited – during study in young adults (Moore & Long, 2024). Thus, increasing numbers of consecutively presented semantically overlapping items could induce young adults to shift attention from the episodic to the semantic dimension during study. As older adults have difficulty inhibiting retrieval of semantic information (Wynn et al., 2020), they may retrieve semantically overlapping items during study, orienting attention toward the semantic dimension of events. This interpretation is consistent with our prior work showing that older adults have a modest bias to engage in memory retrieval during study (Moore, Smith, & Long, 2025). Behaviorally, attending to semantic information during study can promote true memory (Jacoby & Wahlheim, 2013; Tullis et al., 2014; McKinley et al., 2019; McKinley & Benjamin, 2020; Antony et al., 2022; Moore & Long, 2024). However, the observed age differences may arise from differential processing during test rather than study. In older adults, memory errors are particularly likely when participants are tasked with recalling specific details of events (Dodson, Bawa, & Krueger, 2007), suggesting that older adults may “miscombine” features from separate events at test (Chalfonte & Johnson, 1996; Naveh-Benjamin, Hussain, Guez, & Bar-On, 2003). Characterizing encoding and retrieval contributions to age-related differences in reliance on semantic versus episodic information is an important avenue for future work.

In conclusion, we show that young and older adults rely on semantic versus episodic information similarly to support false memory but differentially to support true memory. Our results suggest that episodic memory deficits in older adults may lead to age-related differences in the baseline orientation of attention toward the semantic or episodic dimensions of events. To the extent that attention to semantic versus temporal features of events extend beyond episodic memory, the differences we observe in the present study may reflect broad age-related changes in cognition. These results advance our understanding of how attending to different aspects of events change over the lifespan.

## Acknowledgments

We thank Hannah Buras and Abigail Turner for assistance with data collection. This work was supported by the National Institute on Aging of the National Institutes of Health under Award Number F31AG081045 (to ILM).

## Notes

**Author Note:** This work was supported by the National Institute on Aging of the National Institutes of Health under Award Number F31AG081045 (to ILM). Portions of these findings were presented at the 2024 meeting of the Society for Neuroscience in Chicago, Illinois, United States and the 2025 meeting of the Cognitive Neuroscience Society in Boston, Massachusetts, United States. We have no conflicts of interest to disclose. The raw, deidentified data and the associated experimental and analysis codes used in this study will be available via the Long Term Memory lab website following publication (https://longtermmemorylab.com). The study design and analyses were preregistered (https://osf.io/tkhu9). Correspondence concerning this article should be addressed to Nicole M. Long, Department of Psychology, University of Virginia.; bluesky: @dorsolateralpfc.bsky.social.

### Competing Interest Statement

The authors have declared no competing interest.

## References

Adnan, A., Beaty, R. E., Lam, J., Spreng, R. N., & Turner, G. R. (2019). Intrinsic default-executive coupling of the creative aging brain. Social Cognitive and Affective Neuroscience, 14, 291–303. doi: 10.1093/scan/nsz013

Amer, T., Anderson, J. A., Campbell, K. L., Hasher, L., & Grady, C. L. (2016). Age differences in the neural correlates of distraction regulation: A network interaction approach. NeuroImage, 139, 231–239. doi: 10.1016/j.neuroimage.2016.06.036

Amer, T., Giovanello, K., Grady, C. L., & Hasher, L. (2018). Age differences in memory for meaningful and arbitrary assocations: a memory retrieval account. Psychology and Aging, 33(1), 74–81. doi: 10.1037/pag0000220

Amer, T., Giovanello, K., Nichol, D., Hasher, L., & Grady, C. L. (2019). Neural correlates of enhanced memory for meaningful associations with age. Cerebral Cortex, 29(11), 4568–4579. doi: 10.1093/cercor/bhy334

Antony, J. W., Romero, A., Vierra, A. H., Luenser, R. S., Hawkins, R. D., & Bennion, K. A. (2022). Semantic relatedness retroactively boosts memory and promotes memory interdependence across episodes. eLife, 11, e72519.

Binder, J. R., Desai, R. H., Graves, W. W., & Conant, L. L. (2009). Where is the semantic system? a critical review and meta-analysis of 120 functional neuroimaging studies. Cerebral Cortex, 19(12), 2767–2796. doi: 10.1093/cercor/bhp055

Bower, G. (1972). Coding processes in human memory. In A. W. Melton & E. Martin (Eds.), (p. 85-123). Washington, DC: V.H. Winston.

Brainerd, C. J., & Reyna, V. F. (1990). Gist is the grist: Fuzzy-trace theory and the new intuitionism. Developmental Review, 10, 3–47.

Brainerd, C. J., & Reyna, V. F. (2002). Fuzzy-trace theory and false memory. Current Directions in Psychological Science, 11(5), 164–170. doi: 10.1111/1467-8721.00192

Castel, A. D. (2005). Memory for grocery prices in younger and older adults: The role of schematic support. Psychology and Aging, 20(4), 718–721. doi: 10.1037/0882-7974.20.4.718

Chalfonte, B. L., & Johnson, M. K. (1996). Feature memory and binding in young and older adults. Memory & Cognition, 24(4), 403–416.

De Brigard, F., Langella, S., Stanley, M., Castel, A., & Giovanello, K. (2019). Age-related differences in recognition in associative memory. *Aging*, Neuropsychology, and Cognition, 27 (2), 289–301.

Devitt, A. L., Addis, D. R., & Schacter, D. L. (2017). Episodic and semantic content of memory and imagination: A multilevel analysis. Memory & Cognition, 45, 1078–1094. doi: 10.3758/s13421-017-0716-1

Dodson, C. S., Bawa, S., & Krueger, L. E. (2007). Aging, metamemory, and high-confidence errors: A misrecollection account. Psychology and Aging, 22(1), 122–133.

Dodson, C. S., & Krueger, L. E. (2006). I misremember it well: Why older adults are unreliable eyewitnesses. Psychological Bulletin & Review, 13, 770–775.

Estes, W. (1955). Statistical theory of spontaneous recovery and regression. The Psychological Review, 62(3), 145–154.

Folville, A., Bahri, M. A., Delhaye, E., Salmon, E., D’Argembeau, A., & Bastin, C. (2020). Age-related differences in the neural correlates of vivid remembering. NeuroImage, 206, 116336. doi: 10.1016/j.neuroimage.2019.116336

Fraundorf, S. H., Hourihan, K. L., Peters, R. A., & Benjamin, A. S. (2019). Aging and recognition memory: A meta-analysis. Psychological Bulletin, 145(4), 339–371.

Friendly, M., Franklin, P. E., Hoffman, D., & Rubin, D. C. (1982). The toronto word pool: Norms for imagery, concreteness, orthographic variables, and grammatical usage for 1,080 words. Behavior Research Methods & Instrumentation, 14(4), 375–399.

Gallo, D., Roberts, M., & Seamon, J. (1997). Remembering words not presented in lists: Can we avoid creating false memories? Psychological Bulletin & Review, 4, 271–276.

Geerlings, L., Saliasi, E., Renken, R., Maurits, N., & Lorist, M. (2014). Flexible connectivity in the aging brain revealed by task modulations. Human Brain Mapping, 35, 3788–3804. doi: 10.1002/hbm.22437

Golomb, J. D., Peelle, J., Addis, K., Kahana, M. J., & Wingfield, A. (2008). Effects of adult aging on utilization of temporal and semantic associations during free and serial recall. Memory & Cognition, 36(5), 947–956. doi: 10.3758/MC.36.5.947

Gutchess, A. H., & Schacter, D. L. (2012). The neural correlates of gist-based true and false recognition. NeuroImage, 59(4), 3418–3426.

Healey, M. K. (2018). Temporal contiguity in incidentally encoded memories. Journal of Memory and Language, 102, 28–40.

Hill, P. F., King, D. R., & Rugg, M. D. (2021). Age differences in retrieval-related reinstatement reflect age-related dedifferentiation at encoding. Cerebral Cortex, 31, 106–122. doi: 10.1093/cercor/bhaa210

Howard, M. W., & Kahana, M. J. (2002). When does semantic similarity help episodic retrieval? Journal of Memory and Language, 46, 85–98.

Jacoby, L. L., & Wahlheim, C. N. (2013). On the importance of looking back: The role of recursive remindings in recency judgments and cued recall. Memory & Cognition, 41, 625–637.

Jacoby, L. L., Wahlheim, C. N., Rhodes, M. G., Daniels, K. A., & Rogers, C. S. (2010). Learning to diminish the effects of proactive interference: Reducing false memory for young and older adults. Memory & Cognition, 38(6), 820–829.

Kahana, M. J. (1996). Associative retrieval processes in free recall. Memory & Cognition, 24(1), 103–109.

Koen, J. D., & Rugg, M. D. (2019). Neural dedifferentiation in the aging brain. Trends in Cognitive Sciences, 23(7), 547–559. doi: 10.1016/j.tics.2019.04.012

Koutstaal, W., & Schacter, D. L. (1997). Gist-based false recognition of pictures in older and younger adults. Journal of Memory and Language, 37 (555-583). doi: 10.1006/jmla.1997.2529

Koutstaal, W., Schacter, D. L., & Brenner, C. (2001). Dual task demands and gist-based false recognition of pictures in younger and older adults. Journal of Memory and Language, 44, 399–426.

Koutstaal, W., Schacter, D. L., Galluccio, L., & Stofer, K. (1999). Reducing gist-based false recognition in older adults: Encoding and retrieval manipulations. Psychology and Aging, 14(2), 220–237. doi: 10.1037/0882-7974.14.2.220

Lalla, A., Tarder-Stoll, H., Hasher, L., & Duncan, K. (2022). Aging shifts the relative contributions of episodic and semantic memory to decision-making. Psychology and Aging, 37 (6), 667–680. doi: 10.1037/pag0000700

Lichtenstein, S., & Fischhoff, B. (1977). Do those who know more also know more about how much they know? Organizational Behavior & Human Performance, 20(159-183).

Long, N. M., & Kahana, M. J. (2017). Modulation of task demands suggests that semantic processing interferes with the formation of episodic associations. Journal of Experimental Psychology: Learning, Memory, and Cognition, 43(2), 167–176. doi: 10.1037/xlm0000300

Long, N. M., & Kahana, M. J. (2019). Hippocampal contributions to serial-order memory. Hippocampus, 29(3), 1–8.

Long, N. M., Sperling, M. R., Worrell, G. A., Davis, K. A., Lucas, T., Lega, B., … Kahana, M. J. (2017). Contextually mediated spontaneous retrieval is specific to the hippocampus. Current Biology, 27, 1–6.

McKinley, G. L., & Benjamin, A. S. (2020). The role of retrieval during study: Evidence of reminding from overt rehearsal. Journal of Memory and Language, 114, 104128.

McKinley, G. L., Ross, B. H., & Benjamin, A. S. (2019). The role of retrieval during study: Evidence of reminding from self-paced study time. Memory & Cognition, 47, 877–892.

Melton, A. W. (1970). The situation with respect to the spacing of repetitions and memory. Journal of Verbal Learning and Memory, 9(596-606).

Moore, I. L., & Long, N. M. (2024). Semantic associations restore neural encoding mechanisms. Learning & Memory, 31(3), a053996.124.

Moore, I. L., Smith, D. E., & Long, N. M. (2025). Mnemonic brain state engagement is diminished in healthy aging. Neurobiology of Aging, 151, 76–88.

Nasreddine, Z. S., Phillips, N. A., Bedirian, V., Charbonneau, S., Whitehead, V., Collin, I., … Chertkow, H. (2005). The montreal cognitive assessment, MoCA: A brief screening tool for mild cognitive impairment. Journal of the American Geriatrics Society, 53, 695–699. doi: 10.1111/j.1532-5415.2005.53221.x

Naveh-Benjamin, M., Hussain, Z., Guez, J., & Bar-On, M. (2003). Adult age differences in episodic memory: Further support for an associative-deficit hypothesis. Journal of Experimental Psychology: Learning, Memory, and Cognition, 29(5), 826–837.

Nelson, D. L., Zhang, N., & McKinney, V. M. (2001). The ties that bind what is known to the recognition of what is new. Journal of Experimental Psychology: Learning, Memory, and Cognition, 27 (5), 1147–1159.

Norman, K. A., & Schacter, D. L. (1997). False recognition in younger and older adults: Exploring the characteristics of illusory memories. Memory & Cognition, 25(6), 838–848. doi: 10.3758/BF03211328

Ofen, N., & Shing, Y. L. (2013). From perception to memory: Changes in memory systems across the lifespan. Neuroscience and Biobehavioral Reviews, 37, 2258–2267. doi: 10.1016/j.neubiorev.2013.04.006

Pauley, C., Karlsson, A., & Sander, M. C. (2024). Early visual cortices reveal interrelated item and category representations in aging. eNeuro, 11(3). doi: 10.1523/ENEURO.0337-23.2023

Polyn, S. M., Norman, K. A., & Kahana, M. J. (2009). A context maintenance and retrieval model of organizational processes in free recall. Psychological Review, 116, 129–156.

Postman, L., & Keppel, G. (1977). Conditions of cumulative proactive inhibition. Journal of Experimental Psychology: General, 106(4), 376–403.

Pressley, M., & Ghatala, E. S. (1988). Delusions about performance on multiple-choice comprehension tests. Reading Research Quarterly, 23(4), 454–464.

Raichle, M. E., MacLeod, A. M., Snyder, A. Z., Powers, W. J., Gusnard, D. A., & Shulman, G. L. (2001). A default mode of brain function. Proc. Natl. Acad. Sci., 98(2), 676–682.

Rieck, J., Rodrigue, K. M., Boylan, M., & Kennedy, K. M. (2017). Age-related reduction of bold modulation to cognitive difficulty predicts poorer task accuracy ad poorer fluid reasoning ability. NeuroImage, 147, 262–271. doi: 10.1016/j.neuroimage.2016.12.022

Roebers, C. (2002). Confidence judgements in children’s and adults’ event recall and suggestibility. Developmental Psychology, 38(6), 1052–1067.

Roediger, H. L., & McDermott, K. B. (1995). Creating false memories: Remembering words not presented in lists. Journal of Experimental Psychology: Learning, Memory, and Cognition, 21(4), 800–814.

Salthouse, T. (2011). Effects of age on time-dependent cognitive change. Psychological Science, 22, 682–688. doi: 10.1177/0956797611404900

Schacter, D. L., Israel, L., & Racine, C. A. (1999). Suppressing false recognition in younger and older adults: The distinctiveness heuristic. Journal of Memory and Language, 40, 1–24.

Schneider, S., & Laurion, S. (1993). Do we know what we’ve learned form listening to the news? Memory & Cognition, 21(2), 198–209.

Sederberg, P. B., Miller, J. F., Howard, M. W., & Kahana, M. J. (2010). The temporal contiguity effect predicts episodic memory performance. Memory & Cognition, 38, 689–699.

Shing, Y. L., Werkle-Bergner, M., Li, S.-C., & Lindenberger, U. (2009). Committing memory errors with high confidence: Older adults do but children don’t. Memory, 17 (2), 169–179.

Spreng, R. N., DuPre, E., Selarka, D., Garcia, J., Gojkovic, S., Mildner, J., & Turner, G. R. (2014). Goal-congruent default network activity facilitates cognitive control. Journal of Neuroscience, 34, 14108–14114. doi: 10.1523/JNEUROSCI.2815-14.2014

Spreng, R. N., & Schacter, D. L. (2012). Default network modulation and large-scale network interactivity in healthy young and old adults. Cerebral Cortex, 22, 2610–2621. doi: 10.1093/cercor/bhr339

Spreng, R. N., & Turner, G. R. (2019). The shifting architecture of cognition and brain function in older adulthood. Perspectives on Psychological Science, 14(4), 523–542. doi: 10.1177/1745691619827511

Srokova, S., Aktas, A. N. Z., Koen, J. D., & Rugg, M. D. (2024). Dissociative effects of age on neural differentiation at the category and item levels. The Journal of Neuroscience, 44(4), 1–12. doi: 10.1523/JNEUROSCI.0959-23.2023

Szpunar, K. K., McDermott, K. B., & Roediger, H. L. (2008). Testing during study insulates against the buildup of proactive interference. Journal of Experimental Psychology: Learning, Memory, and Cognition, 34(6), 1392–1399.

Tullis, J., Benjamin, A. S., & Ross, B. H. (2014). The reminding effect: Presentation of associates enhances memory for related words in a list. Journal of Experimental Psychology: General, 143(4), 1526–1540.

Tulving, E. (1972). Episodic and semantic memory. Organization of Memory. Academic Press.

Tun, P. A., Wingfield, A., Rosen, M. J., & Blanchard, L. (1998). Response latencies for false memories: Gist-based processes in normal aging. Psychology and Aging, 13(2), 230–241. doi: 10.1037//0882-7974.13.2.230

Turner, G. R., & Spreng, R. N. (2015). Prefrontal engagement and reduced default network suppression co-occur and are dynamically coupled in older adults: The default-executive coupling hypothesis of aging. Journal of Cognitive Neuroscience, 27, 2462–2476. doi: 10.1162/jocna00869

Underwood, B. J. (1957). Interference and forgetting. Psychological Review, 64(1), 49–60.

Unsworth, N., Brewer, G. A., & Spillers, G. J. (2013). Focusing the search: Proactive and retroactive interference and the dynamics of free recall. Journal of Experimental Psychology: Learning, Memory & Cognition, 39, 1742–1756.

Wahlheim, C. N., & Huff, M. J. (2015). Age differences in the focus of retrieval: Evidence from dual-list free recall. Psychology and Aging, 30(4), 768–780.

Wang, F., & Diana, R. A. (2017). Temporal context in human fmri. Current Opinion in Behavioral Sciences, 17, 57–64.

Wilkniss, S., Jones, M., Korol, D., Gold, P., & Manning, C. (1997). Age-related differences in an ecologically based study of route learning. Psychology and Aging, 12(2), 372–375. doi: 10.1037/0882-7974.12.2.372

Wingfield, A., & Kahana, M. J. (2002). The dynamics of memory retrieval in older adulthood. Canadian Journal of Experiemental Psychology, 56(3), 187–199. doi: 10.1037/h0087396

Wingfield, A., Lindfield, K. C., & Kahana, M. J. (1998). Adult age differences in the temporal characteristics of category free recall. Psychology and Aging, 13(2), 256–266.

Wynn, J., Ryan, J. D., & Moscovitch, M. (2020). Effects of prior knowledge on active vision and memory in younger and older adults. Journal of Experimental Psychology: General, 149(3), 518–529. doi: 10.1037/xge0000657

Yonelinas, A., Ranganath, C., Ekstrom, A., & Wiltgen, B. (2019). A contextual binding theory of episodic memory: Systems consolidation reconsidered. Nature Reviews Neuroscience, 20(6), 364–375.

